# Neuroligin3 Splice Isoforms Shape Mouse Hippocampal Inhibitory Synaptic Function

**DOI:** 10.1101/2020.01.22.915801

**Authors:** Motokazu Uchigashima, Ming Leung, Takuya Watanabe, Amy Cheung, Masahiko Watanabe, Yuka Imamura Kawasawa, Kensuke Futai

## Abstract

Synapse formation is a dynamic process essential for neuronal circuit development and maturation. At the synaptic cleft, trans-synaptic protein-protein interactions constitute major biological determinants of proper synapse efficacy. The balance of excitatory and inhibitory synaptic transmission (E-I balance) stabilizes synaptic activity. Dysregulation of the E-I balance has been implicated in neurodevelopmental disorders including autism spectrum disorders. However, the molecular mechanisms underlying E-I balance remain to be elucidated. Here, we investigate Neuroligin (*Nlgn*) genes that encode a family of postsynaptic adhesion molecules known to shape excitatory and inhibitory synaptic function. We demonstrate that Nlgn3 protein differentially regulates inhibitory synaptic transmission in a splice isoform-dependent manner at hippocampal CA1 synapses. Distinct subcellular localization patterns of Nlgn3 isoforms contribute to the functional differences observed among splice variants. Finally, single-cell sequencing analysis reveals that *Nlgn1* and *Nlgn3* are the major *Nlgn* genes and that expression of *Nlgn* splice isoforms are highly diverse in CA1 pyramidal neurons.

## INTRODUCTION

Neuroligin proteins (Nlgns) were the first identified binding partners of α-latrotoxin receptors, neurexin proteins (Nrx), and localize at postsynaptic sites to regulate synapse maturation and function (1). Four Nlgn genes (*Nlgn1, Nlgn2, Nlgn3* and *Nlgn4*) encode trans-synaptic adhesion proteins (Nlgn1, Nlgn2, Nlgn3 and Nlgn4) that contain extracellular cholinesterase-like domains and transmembrane and PDZ-binding motif-containing intracellular domains. While the intracellular domain is important for NLGN binding with postsynaptic scaffold molecules, the extracellular domain confers the formation of excitatory and inhibitory synapses with Nrx, its sole presynaptic binding partner. Therefore, precise combinations of Nrx-Nlgn interactions allow Nlgns to diversify synapse identity.

Nlgn1 and Nlgn2 are postsynaptic adhesion molecules localized to excitatory and inhibitory synapses, respectively. Overexpression, knockdown and knockout approaches have revealed that Nlgn1 is important for excitatory synaptic structure and transmission and synaptic plasticity, but not for inhibitory synaptic function (2–6). Nlgn2 has specific functional roles in inhibitory synaptic transmission in the hippocampus (6–8). In contrast, it has been reported that Nlgn3 protein localizes at both excitatory and inhibitory synaptic sites and regulates both synaptic functions (7–12). This unique ability alludes to a Nlgn3 protein-specific molecular code that promotes its translocation to both excitatory and inhibitory sites.

Splice insertion in *Nlgn* genes differentially regulates E-I balance and alters their binding affinity with presynaptic Nrx partners. In the extracellular cholinesterase-like domain, *Nlgn* genes have one or two splice insertion sites, *Nlgn1* (A and B sites), *Nlgn2* (A) and *Nlgn3* (A1 and A2), leading to two to four theoretical splice isoforms. The splice insertion site B in Nlgn1 determines its binding preference to Nrxs (13, 14) and excitatory synaptic function (15). Similarly, Nlgn2 contains a splice insertion at site A which regulates inhibitory synaptic function (8). However, to the best of our knowledge, the splice isoform-specific function of Nlgn3 and the transcript levels of Nlgn splice isoforms at the single-cell level have not been addressed.

In the present study, we assess the function of Nlgn3 splice isoforms on excitatory and inhibitory synaptic transmission in CA1 pyramidal neurons in mouse organotypic slice cultures. Our results suggest that Nlgn3 up- or down-regulates inhibitory synaptic transmission in a splice isoform-dependent manner. Furthermore, our single-cell RNA sequencing (RNA-seq) analysis reveals that *Nlgn1* and *Nlgn3* are the major *Nlgn* genes and the expression of *Nlgn* splice variants are highly distinct in hippocampal CA1 pyramidal neurons.

## Results

### Nlgn3 splice isoform-dependent regulation of inhibitory synaptic transmission

The Nlgn3 gene contains two splice insertion sites (A1 and A2) that can yield four Nlgn3 splice isoforms (Nlgn3Δ, A1, A2 and A1A2). Nlgn3Δ lacks all splice insertions, while Nlgn3A1, 3A2 and 3A1A2 receive insertion of A1, A2 or both A1A2 cassette(s), respectively. To examine the potential roles of Nlgn3 splice isoforms on excitatory and inhibitory synapse function, we biolistically transfected the Nlgn3 splice isoforms in CA1 pyramidal cells of organotypic hippocampal slice cultures (**Fig. 1**). Simultaneous electrophysiological recordings were made from transfected and neighboring untransfected neurons. CA1 pyramidal neurons overexpressing Nlgn3Δ or 3A2 showed increased evoked IPSCs compared with neighboring untransfected control neurons and a marked increase in EPSCs, as reported previously (**Fig. 1A and C**) (7, 12). In contrast, overexpression of Nlgn3A1 or 3A1A2 resulted in reduced amplitude of IPSCs and increased amplitude of EPSCs compared with neighboring untransfected cells (**Fig. 1B and D**). Paired stimulation of input fibers with short interval (50 ms) induced paired-pulse facilitation (PPF) and depression (PPD) of EPSCs and IPSCs, respectively. Nlgn3Δ or 3A2 transfection displayed both reduced AMPAR-PPF and GABA_A_R-PPD compared with untransfected neurons, consistent with previous work (5, 7) (**Fig. S1B and C**). As paired-pulse ratio (PPR) inversely correlates with presynaptic release probability, these results suggest that overexpression of Nlgn3Δ and 3A2 can modulate presynaptic release probability. Nlgn3 increased or decreased inhibitory synaptic transmission in a splice isoform-dependent manner, whereas all Nlgn3 splice isoforms enhanced excitatory synaptic transmission.

**Figure 1.**
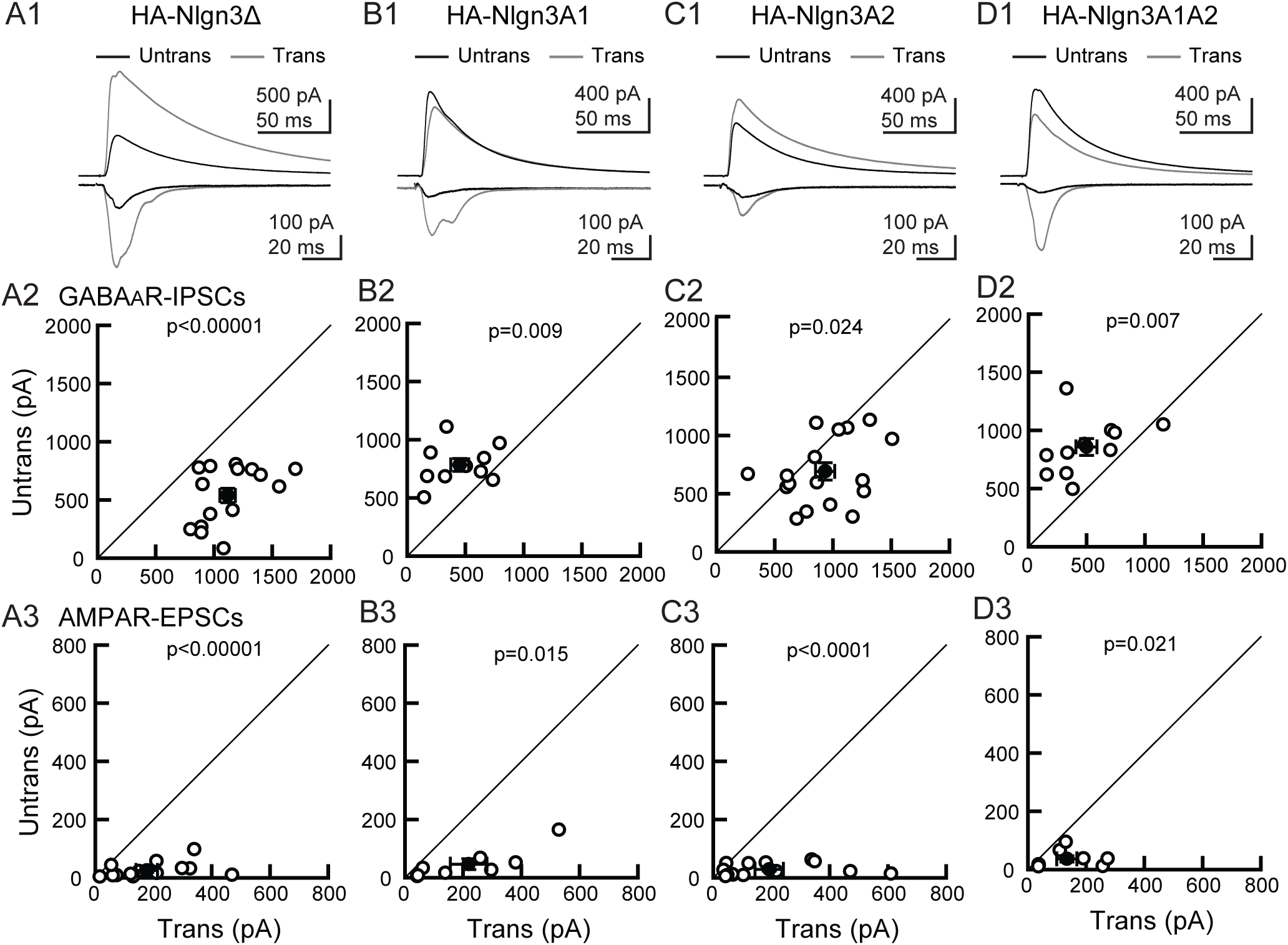
Nlgn3 splice isoforms differentially encode synapse specificity. Effect of Nlgn3 splice isoform overexpression on GABA_A_R-IPSCs and AMPAR-EPSCs in hippocampal CA1 pyramidal cells. (A1-D1) Top row: superimposed sample IPSCs and EPSCs from pairs of transfected (gray) and untransfected (black) cells. Stimulus artifacts were truncated. (A) Nlgn3Δ. (B) 3A1. (C) 3A2. (D) 3A1A2. IPSC (middle row, A2-D2) and EPSC (bottom row, A3-D3) amplitudes were plotted for each pair of transfected (Trans) and neighboring untransfected (Untrans) cells (open symbols). Filled symbols indicate the mean ± s.e.m. Numbers of cell pairs: Nlgn3Δ (IPSCs/ EPSCs: 16/14); 3A1 (10/8); 3A2 (16/14); 3A1A2 (11/8). Number of tested slice cultures are the same as that of cell pairs. Mann–Whitney U test.

### Subcellular localization of Nlgn3 splice isoforms in the dendritic segment of CA1 pyramidal neurons

Expression of Nlgn3 at excitatory and inhibitory synapses has been observed in primary neurons but *in vivo* Nlgn3 expression has been studied only in the cerebellum and striatum (11,16,17). To ensure the expression of Nlgn3 in the hippocampus, we performed immunohistochemistry against Nlgn3 with the markers for excitatory and inhibitory synapses, VGluT1 and VIAAT, respectively (**Fig. 2**). Our Nlgn3 antibody, validated by Nlgn3 knockout tissue, detected punctate signals in the hippocampus (**Fig. 2A and B**). The signals overlapped with VGluT1 and VIAAT puncta, indicating that Nlgn3 proteins are targeted to both excitatory and inhibitory synapses, respectively (**Fig. 2C and D**). To understand the mechanistic roles of Nlgn3 splice isoforms in inhibitory synaptic transmission, we next performed immunocytochemistry to elucidate the subcellular localization of Nlgn3Δ and 3A1A2, which displayed strong enhancement and suppression of IPSC, respectively. Excitatory synaptic sites were characterized by spine or VGluT1. Inhibitory synaptic sites were identified by the dendritic shaft proximal to VIAAT puncta. HA immunoreactivity illustrated that Nlgn3A1A2 is highly concentrated in spines. In contrast, Nlgn3Δ showed more diffuse expression in both spines and dendrites (**Fig. 3A**). The ratio of Nlgn3A1A2 signals between excitatory and inhibitory synapses was significantly higher than that of 3Δ (**Fig. 3B**). Next, we addressed whether these Nlgn3 splice isoforms differentially promote excitatory and inhibitory synapses. Importantly, inhibitory synapse density was comparable between Nlgn3Δ and 3A1A2, whereas VIAAT signal intensity in 3A1A2-expressing neurons was significantly lower than that of 3Δ (**Fig. 3C and D**). The spine density was comparable between Nlgn3Δ- and 3A1A2-overexpressing neurons (**Fig. 3E**). The signal intensities of VGluT1 were markedly elevated in neurons overexpressing Nlgn3Δ or 3A1A2 compared with the surrounding VGluT1 puncta representing excitatory synaptic sites of untransfected neurons (**Fig. 3F and G**). These results suggest that differences in the subcellular localization of Nlgn3Δ and 3A1A2 contribute to their distinct inhibitory synaptic functions.

**Figure 2.**
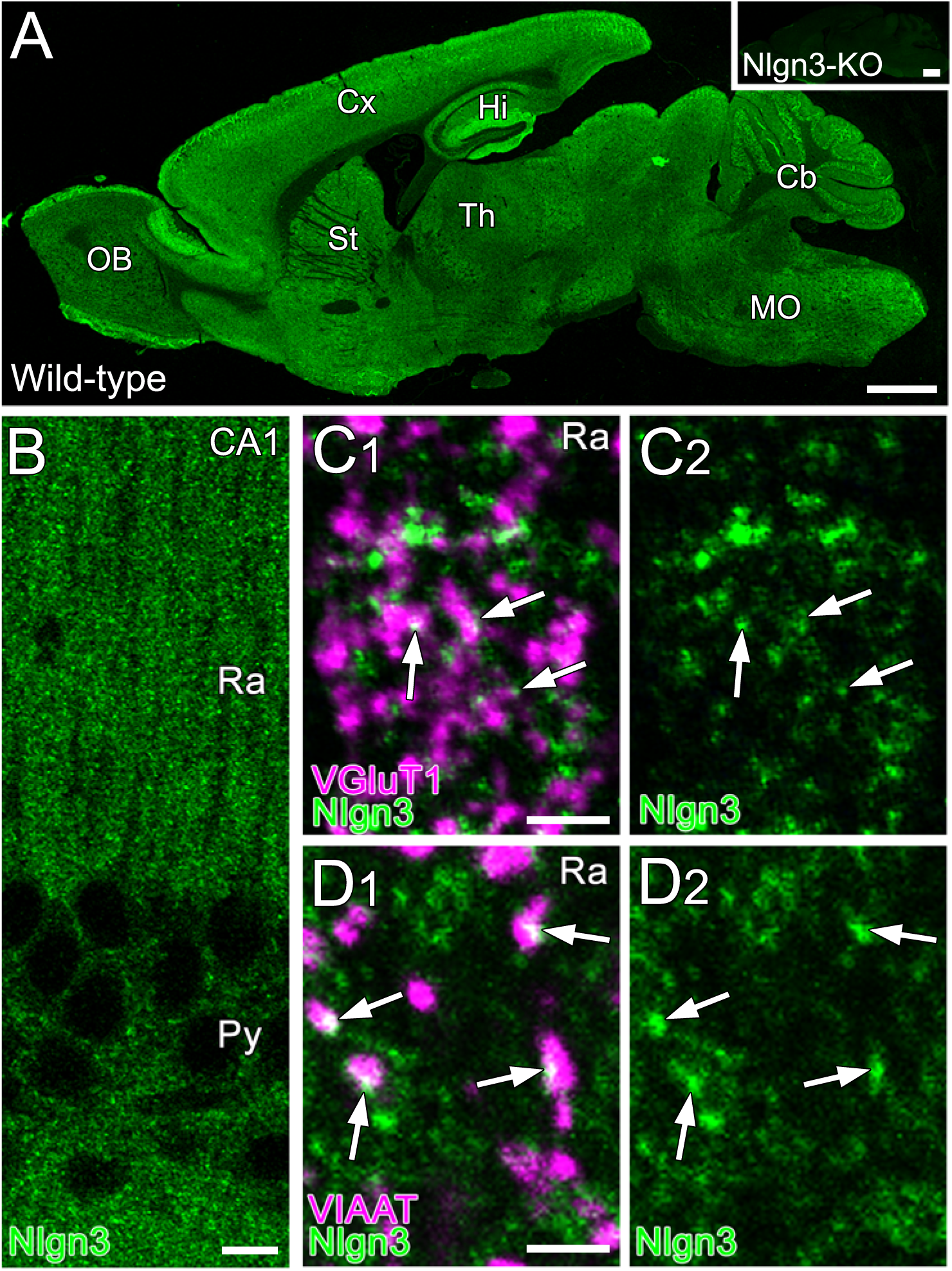
Subcellular localization of Nlgn3 in the hippocampal CA1 area. (A) Immunohistochemical staining for Nlgn3 in whole brains of wild-type and Nlgn3-knockout mice (Nlgn3-KO, inset). (B) Immunohistochemical staining for Nlgn3 in the hippocampal CA1 area. (C, D) Double immunohistochemical staining for Nlgn3 (green) and VGluT1 (C, magenta) or VIAAT (D, magenta) in the stratum radiatum of the hippocampal CA1 area. Cb, cerebellum; Cx, cortex; Hi, hippocampus; MO, medulla oblongata; OB, olfactory bulb; Py, stratum pyramidale; Ra, stratum radiatum; St, striatum; Th, thalamus. Scale bars: 1 mm (A), 10 µm (B), 2 µm (C, D).

**Figure 3.**
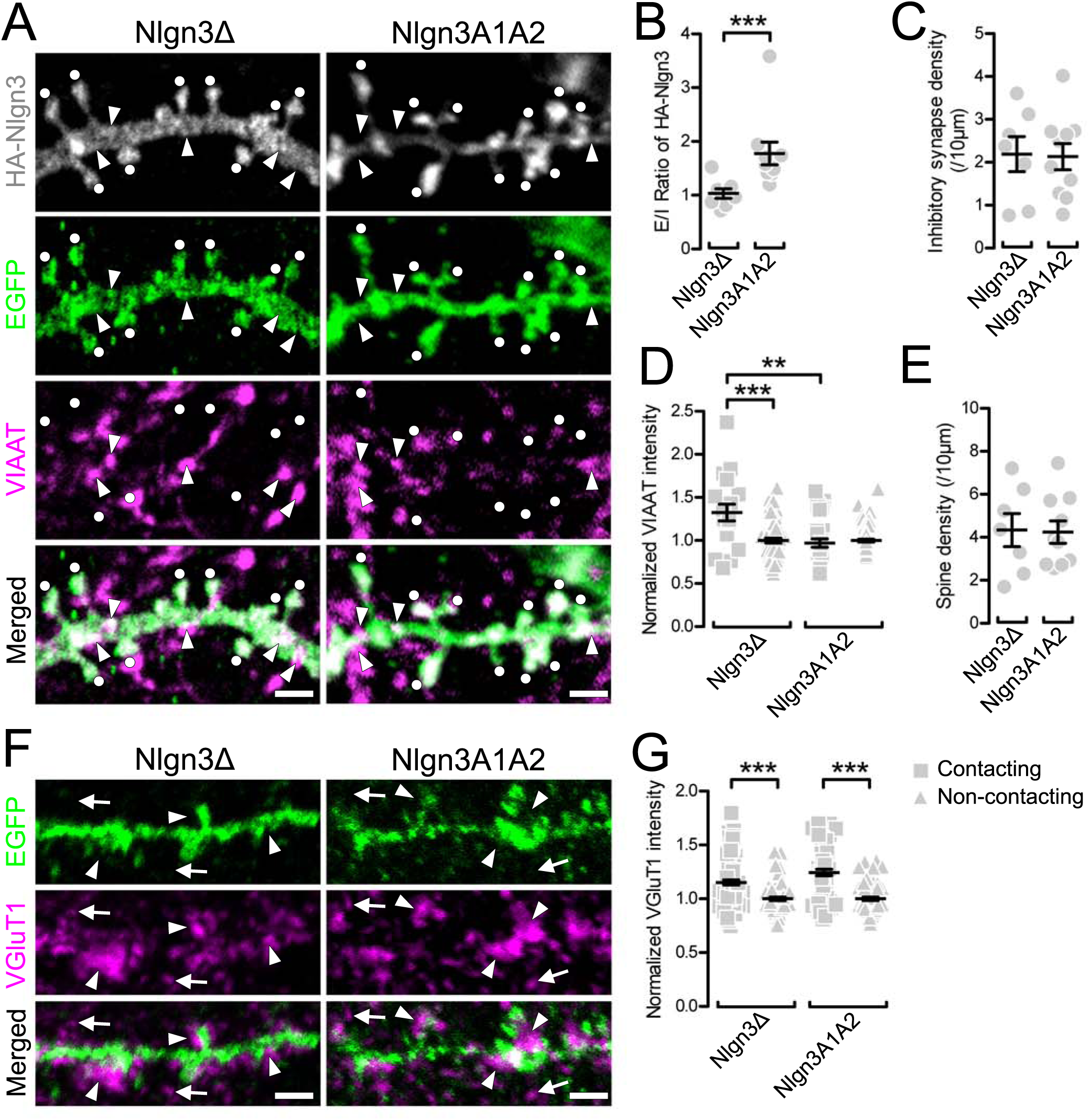
Synaptic targeting of Nlgn3 splice isoforms in CA1 pyramidal neurons. (A) Maximum projection images of dendritic segments labeled for HA-Nlgn3 (gray), EGFP (green) and VIAAT (magenta) in CA1 pyramidal cells overexpressing HA-Nlgn3Δ (left) and A1A2 (right). White dots and arrows indicate EGFP+ spines and VIAAT+ inhibitory synapses, respectively. Scale bars: 2 µm. (B-D) Summary scatter plot showing the E/I ratio of HA signals (B), density of inhibitory synapses(C), normalized VIAAT intensity (D), and spine density (E) for individual dendritic segments of CA1 pyramidal cells overexpressing HA-Nlgn3Δ (left, n = 7 dendrites) and A1A2 (right, n = 10). Normalized VIAAT intensity is obtained from contacting (square) and non-contacting (triangle) terminals to HA-Nlgn3-overexpressing dendrites. (F) Maximum projection images of dendritic segments labeled for EGFP (green) and VGluT1 (magenta) in CA1 pyramidal cells overexpressing HA-Nlgn3Δ (left) and A1A2 (right). Arrowheads and arrows indicate VGluT1-labeled terminals contacting and not contacting to dendrites overexpressing HA-Nlgn3, respectively. (G) Summary scatter plot showing the normalized VGluT1 intensity in contacting (square) and non-contacting (triangle) terminals to individual dendritic segments of CA1 pyramidal cells overexpressing HA-Nlgn3Δ (left, n = 7 dendrites) and A1A2 (right, n = 6). Scale bars: 2 µm. ** p < 0.01, *** p < 0.001 (one-way ANOVA followed by Sidak’s multiple comparisons test or U-test).

### Endogenous expression of *Nlgn* genes and splice isoforms in hippocampal CA1 pyramidal neurons

Finally, to understand the expression of endogenous *Nlgn* genes in CA1 pyramidal neurons, we harvested cytosol from four neurons and performed single-cell deep RNA-seq. The t-SNE plot indicates that the four cell transcripts (G418) were clustered together and close to that of adult hippocampal CA1-3 pyramidal neurons derived from the Allen Brain Atlas (**Fig. 4A**). The expression of Nlgn genes was clustered and well correlated with the single-cell RNA-seq data in the RNA-seq datasets provided by the Allen Institute for Brain Science (**Fig. 4B**). The quantification of *Nlgn* genes (**Fig. 4C**) indicates that the expression of *Nlgn1* and *Nlgn3* are comparable but that of *Nlgn2* is significantly lower than the other two genes. We also compared the expression of *Nlgn* splice isoforms in each *Nlgn* gene. Six *Nlgn* splice isoforms, *Nlgn1A, 1B, 2Δ, 3Δ* and *3A1*, that were not annotated, were manually modified (**Fig. S2**), and their expression was compared. *Nlgn1Δ, 1B* and *1AB* were the most highly expressed Nlgn1 splice isoforms (**Fig. 4D**). *Nlgn2Δ* was the only isoform counted in the *Nlgn2* gene (**Fig. 4E**). *Nlgn1A* and *2A* transcripts were not detected in any of the four CA1 pyramidal neurons. Importantly, *Nlgn3Δ* and *3A2*, which exhibited increased inhibitory synaptic transmission, were the dominant *Nlgn3* splice isoforms in CA1 pyramidal neurons (**Fig. 4F**). The expression of *Nlgn3A1* and *3A1A2* were significantly lower than *Nlgn3Δ* (TPM: *Nlgn3A1*: 0.004 ± 0.004, *3A1A2*: 0.3 ± 0.3).

**Figure 4.**
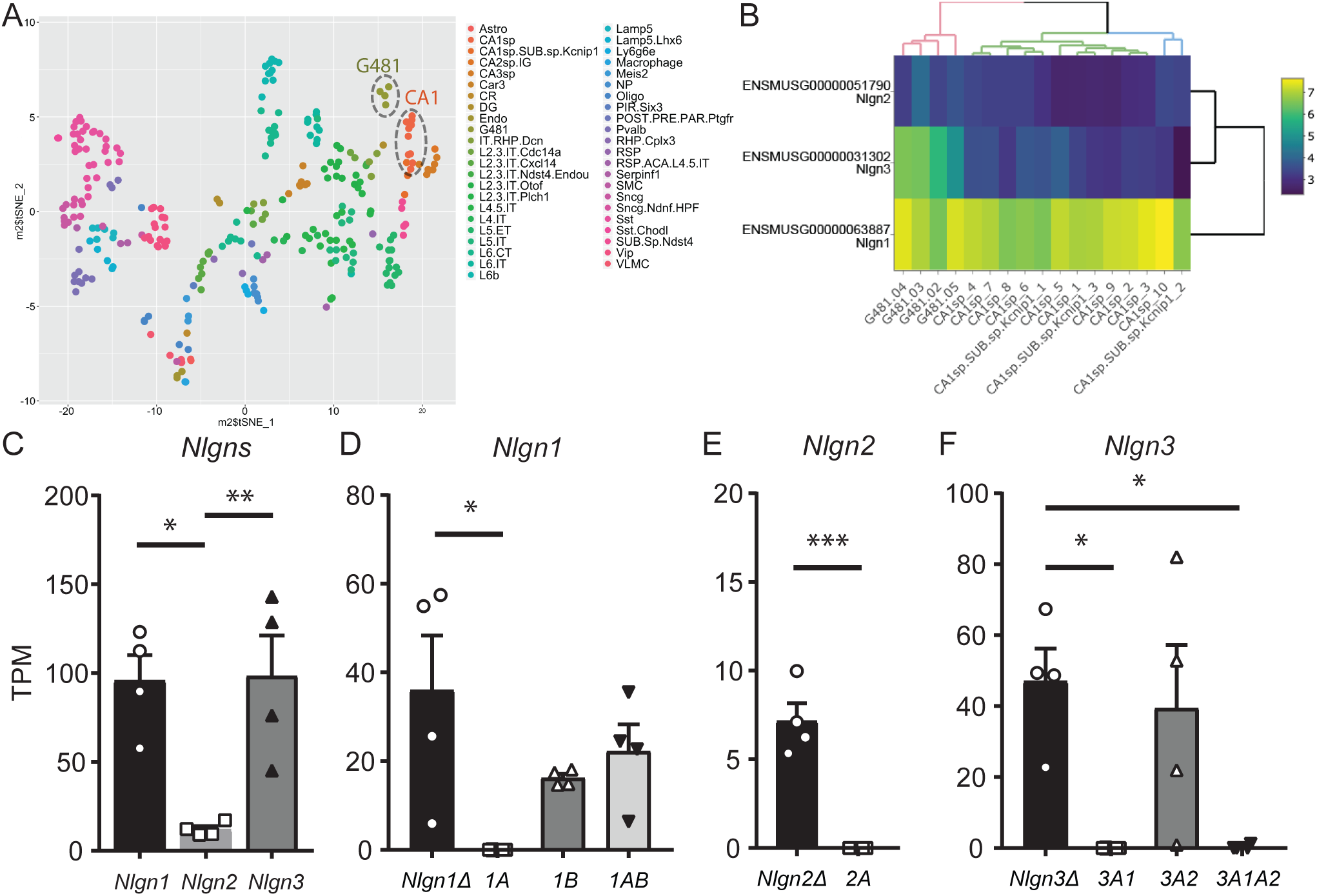
Endogenous *Nlgn* expression in hippocampal CA1 pyramidal neurons. (A) Single-cell t-SNE plot of four single-cells (G481.02, .03, .04 and .05) and Allen Brain Atlas single-cells. (B) Heat map of *Nlgn* gene expression in hippocampal CA1 primary neurons. A hierarchical clustering demonstrated a segregation of G481 cells from adult CA1 pyramidal neurons for *Nlgn3*, potentially due to relatively higher expression of *Nlgn3* in G481 cells. Scale bar shows log2+1 transformed TPM values. (C) Summary bar graph of Nlgn gene (*Nlgn1, 2* and *3*) expression. (D-F) Summary graphs of splice isoforms of *Nlgn1* (D), *2* (E) and *3* (F). * p < 0.05, one-way ANOVA followed by Sidak’s multiple comparisons test or U-test, n = 4.

## Discussion

Trans-synaptic protein-protein interactions are fundamental biological events for synapse formation, maturation and function. Nlgns are critical postsynaptic adhesion molecules that regulate excitatory and inhibitory synaptic transmission. Here we demonstrated that Nlgn3 regulates inhibitory synaptic transmission and excitatory and inhibitory synapse localization in a splice isoform-dependent manner. Our single-cell transcriptome analysis revealed that *Nlgn3Δ* and *3A2* are the highest expressed *Nlgn3* splice isoforms in hippocampal CA1 pyramidal neurons.

The distinct subcellular localization of Nlgn3Δ and 3A1A2 suggests intriguing mechanisms regarding how splice isoforms influence synapse specificity. Given that the intracellular and transmembrane domains are identical between Nlgn3 splice isoforms, each isoform exerts their synapse coding effect through their unique extracellular domains. Similarly, the extracellular domain of Nlgn2 mediates changes in inhibitory synaptic function (6). Although cis-protein interactions with Nlgn3s cannot be excluded (18), trans-synaptic interactions between Nlgn3Δ, but not 3A1A2, and Nrxs modulate inhibitory synaptic transmission in pyramidal neurons. We previously reported that postsynaptic Nlgn2 can couple with presynaptic αNrx1 but not with βNrx1 to form functional inhibitory synapses (6), suggesting that the specific binding of Nlgn and Nrx isoforms regulates functional synapse formation. It is possible that inhibitory interneurons do not express Nrx isoforms that can bind to Nlgn3A1A2. Therefore, Nlgn3A1A2 may exclusively localize to excitatory synapses. Further studies should be performed to identify specific Nrx-Nlgn3 isoform interactions which affect inhibitory synaptic function. It has been suggested that the relative levels of Nlgns and their postsynaptic scaffold complex at excitatory and inhibitory synapses determine E-I balance (19). Additional studies are required, but it is interesting to address the hypothesis that postsynaptic Nlgn3A1 and A1A2 are strong and specific regulators at excitatory synaptic sites and sequester the necessary protein interactions (i.e., Nlgn2-mediated scaffold complex) from inhibitory synapses to reduce inhibitory synaptic transmission.

The single-cell sequencing results demonstrate a comprehensive unbiased gene expression profile of *Nlgn* splice isoforms in hippocampal CA1 pyramidal neurons obtained from neonatal mice. Our transcriptome findings are highly correlated with the expression pattern of the three *Nlgns* genes provided by the Allen Brain Atlas single-cell database from adult neurons, indicating that the expression ratio of *Nlgns* are stable throughout development. Interestingly, Nlgn2 has been well-characterized at inhibitory synapses, yet its transcript levels were significantly lower than *Nlgn1* and *Nlgn3* (**Fig. 4B and C**). Nlgn2 may have unique post-translational modifications and turnover mechanisms compared with Nlgn1 and Nlgn3. The expression of *Nlgn3A1* and *3A1A2* were much lower than *Nlgn3Δ* and *3A2*, suggesting that these two splice isoforms are not the major functional *Nlgn3* molecules under basal conditions. Further work is necessary to address whether modifications to neuronal activity such as synaptic plasticity can alter the expression of *Nlgn* splice isoforms.

A shift in E-I balance has been considered a pathophysiological hallmark of neurodevelopmental disorders and repeatedly reported in corresponding mouse models (20). Mutations and deletions of *Nlgn3* loci are associated with autism spectrum disorders (21), and mutant mice that mimic the human autism *Nlgn3* mutation exhibit E-I imbalance and abnormal synaptic plasticity (1). A closer examination of specific Nlgn3 splice isoform functions will elucidate their role in producing critical molecular outcomes that may influence neuropsychiatric disease pathogenesis.

## Experimental procedures

### Single-cell sequencing and analysis

*Single-Cell RNA Extraction:* The cytosol of four CA1 neurons were harvested using the whole cell patch-clamp technique described previously (6). Briefly, glass electrodes (1.5 – 2.0 MΩ) were filled with DEPC-treated internal solution containing (in mM): 140 K-methanesulfonate, 0.2 EGTA, 2 MgCl_2_ and 10 HEPES, pH-adjusted to 7.3 with KOH. RNase inhibitor (1U/μl, Ambion) was included in the internal solution. Immediately after establishing whole-cell mode, the cytosol of the recorded cell was aspirated into the patch pipette and immediately expelled into an RNase-free 0.5-ml tube (Ambion). *Library Preparation and mRNA Sequencing:* The cDNA libraries were prepared using a SMART-Seq® HT Kit (TAKARA Bio) and a Nextera XT DNA Library Prep Kit (Illumina) as per the manufacturers’ instructions. Unique barcode sequences were incorporated into the adaptors for multiplexed high-throughput sequencing. The final product was assessed for its size distribution and concentration using a BioAnalyzer High Sensitivity DNA Kit (Agilent Technologies). The libraries were pooled and diluted to 3 nM using 10 mM Tris-HCl (pH 8.5) and then denatured using the Illumina protocol. The denatured libraries were loaded onto an S1 flow cell on an Illumina NovaSeq 6000 (Illumina) and run for 2 x 50 cycles according to the manufacturer’s instructions. De-multiplexed sequencing reads were generated using Illumina bcl2fastq (released version 2.18.0.12) allowing no mismatches in the index read. *Data Analysis*: BBDuk (https://jgi.doe.gov/data-and-tools/bbtools/bb-tools-user-guide/bbduk-guide/) was used to trim/filter low quality sequences using the “qtrim=lr trimq=10 maq=10” option. Next, alignment of the filtered reads to the mouse reference genome (GRCm38) was performed using HISAT2 (version 2.1.0) (https://genomebiology.biomedcentral.com/articles/10.1186/gb-2013-14-4-r36) applying --no-mixed and --no-discordant options. Read counts were calculated using HTSeq (http://bioinformatics.oxfordjournals.org/content/31/2/166) by supplementing Ensembl gene annotation (GRCm38.78). Gene expression values were calculated as transcripts per million (TPM) using custom R scripts. Genes with no detected TPM in all samples were filtered out. The log2+1 transformed TPM values were combined with “Cell Diversity in the Mouse Cortex and Hippocampus RNA-Seq Data” from the Allen Institute for Brain Science (https://portal.brain-map.org/atlases-and-data/rnaseq#Mouse_Cortex_and_Hip) where large-scale single-cell RNA-seq data was collected from adult mouse brain. RNA sequencing data were generated from single cells isolated from >20 areas of mouse cortex and hippocampus, including ACA, AI, AUD, CA, CLA, CLA;EPd, ENTl, ENTm, GU;VISC;AIp, HIP, MOp, MOs, ORB, PAR;POST;PRE, PL;ILA, PTLp, RSP, RSPv, SSp, SSs, SSs;GU, SSs;GU;VISC, SUB;ProS, TEa;PERI;ECT, VISal;VISl;VISli, VISam;VISpm, VISp and VISpl;VISpor (Abbreviations for each cell type match those found in the Allen Mouse Brain Atlas (https://portal.brain-map.org/explore/classes/nomenclature)). The provided table of median expression values for each gene in each cell type cluster was merged with our data, and a tSNE plot was generated using Rtsne R package (22). For splice isoform quantification, kallisto (23) was used by supplementing the transcript fasta file (Mus_musculus.GRCm38.cdna.all.fa), which was manually modified to include six *Nlgn* splice isoforms (*Nlgn1A*_XM_006535423.3, *1B*_XM_006535424.4, *2Δ*_XM_006532902.4, *2A*_XM_006532903.3, *3Δ* and *3A1*) that were not annotated by the Mus_musculus.GRCm38.cdna.all.fa dataset. The manually curated transcript sequences are provided in **Fig. S2**.

### Immunohistochemistry and immunocytochemistry

Mice were transcardially perfused with 4% paraformaldehyde/0.1 M phosphate buffer. Brains were dehydrated and embedded with paraffin. Paraffin sections (2 µm in thickness) were generated using a sliding microtome (Leica). Prior to immunoreactions, paraffin sections were boiled with Immunosaver (Nisshin EM) for 30 min. Organotypic slice cultures transfected with Nlgn3 splice isoforms were fixed with 4% PFA and 4% sucrose in 0.01 M phosphate-buffered saline (PBS).

Organotypic slices were permeabilized with 0.1-0.5% Triton X-100 / PBS (PBST), followed by blocking with 10% goat serum. A mixture of primary antibodies against enhanced green fluorescent protein (EGFP: chicken, Millipore), human influenza hemagglutinin (HA; rabbit, Cell signals), Nlgn3 (guinea pig, (16)), vesicular glutamate transporter type 1 (VGluT1; rabbit and guinea pig, (24)) and vesicular inhibitory amino acid transporter (VIAAT; guinea pig and goat, (25)), and a mixture of species-specific secondary antibodies conjugated with Alexa 488, 555 and 647 (Thermo Fisher) were used for immunostaining. Images were taken with a confocal microscope (FV1200, Olympus) equipped with a 60x silicone oil immersion objective (UPLSAPO 60XS) and analyzed with ImageJ software.

### Electrophysiology

The extracellular solution consisted of (in mM) 119 NaCl, 2.5 KCl, 4 CaCl_2_, 4 MgCl_2_, 26 NaHCO_3_, 1 NaH_2_PO4, 11 glucose and 0.01 2-chloroadenosine (Sigma) gassed with 5% CO_2_ and 95% O_2_, pH of 7.4. Thick-walled borosilicate glass pipettes were pulled to a resistance of 2.5 – 4.0 MΩ. Neurons at DIV 4-6 were transfected using a biolistic gene gun (Bio-Rad) and were assayed 3 days after transfection as described previously (5,6,26,27). Whole-cell voltage clamp recordings were performed with internal solution containing (in mM): 115 cesium methanesulfonate, 20 CsCl, 10 HEPES, 2.5 MgCl_2_, 4 ATP disodium salt, 0.4 guanosine triphosphate trisodium salt, 10 sodium phosphocreatine and 0.6 EGTA, at pH 7.25 adjusted with CsOH. GABA_A_ receptor-mediated inhibitory postsynaptic currents (IPSCs) were measured at *V*hold ± 0 mV. AMPAR-mediated excitatory postsynaptic currents (EPSCs) were evoked at *V*hold -70 mV in the presence of picrotoxin (0.1 mM, Sigma). Recordings were performed using a MultiClamp 700B amplifier and Digidata 1440 and digitized at 10 kHz and filtered at 4 kHz by low-pass filter. Data were acquired and analyzed using pClamp (Molecular Devices).

### Statistical analyses

Results are reported as mean ± s.e.m. Mann-Whitney U-test and Student’s t-test were used for two-group comparison. Statistical significance was set at p < 0.05 for Student’s t-test and U-test.

Other experimental procedures are described in the **supporting Experimental procedures**.

## Author contributions

M.U., T.W., Y.I.K. and K.F. designed research; M.U., M.L., T.W., Y.I.K. and K.F. carried out experiments; M.U., M.L., T.W., Y.I.K. and K.F. analyzed data; M.U., T.W., A.C., M.W., Y.I.K. and K.F. wrote the paper.

## Acknowledgement

This work was supported by the grants from the National Institutes of Health Grants (R01NS085215 to K.F., T32 GM107000 to A.C.), Grants-in-Aid for Scientific Research (15K06732 to M.U.). The authors thank Ms. Naoe Watanabe for skillful technical assistance. We thank Dr. Paul D. Gardner for comments on the manuscript.

## Supporting Experimental Procedure

### Animal and organotypic slice culture preparation

All animal protocols were approved by the Institutional Animal Care and Use Committee (IACUC) at the University of Massachusetts Medical School and the Hokkaido University. Nlgn3 KO mouse line was gift from Dr. Tanaka (1). Organotypic hippocampal slice cultures were prepared from postnatal 6- to 7-day-old C57BL6 mice of both sexes as described previously (2).

### Plasmid constructs

The expression vector for HA-tagged mouse Nlgn3Δ has been reported previously (3). Three HA-tagged Nlgn3 splice variants, Nlgn3A1, Nlgn3A2 and Nlgn3A1A2, were cloned using a PCR-based method.

### Analysis of immunocytochemistry images

For quantification of HA signals, the peak signal intensity was measured at a pair of excitatory and inhibitory synapses on dendritic segments which were identified as a GFP-labeled spine and its neighboring VIAAT-labeled inhibitory terminal, respectively. E-I balance ratio was calculated by dividing the peak signal intensity at excitatory synapses by that at inhibitory synapses. The background signal was determined as the mean signal intensity in GFP-unlabeled regions, and then subtracted from the raw value for the peak signal intensity. To assess the labeling intensity for VIAAT, the contour of VIAAT+ terminals was determined using binary images for the VIAAT channel. The mean intensity was measured in VIAAT+ terminals contacting and not contacting to Nlgn3-overexpressing dendrites, and then normalized with the average intensity for VIAAT+ terminals which were not in contact with Nlgn3-overexpressing dendrites on the same image. This measurement was also applied to the labeling intensity for VGluT1. To assess the density of excitatory and inhibitory synapses, the number of spines and VIAAT+ inhibitory synapses were counted on individual dendritic segments, respectively. The mean length of dendritic segments was 19.7 ± 3.8 µm and 21 ± 2.1 µm for CA1 pyramidal cells overexpressing HA-Nlgn3Δ and 3A1A2, respectively.

### DATA availability

The accession number for the RNA-seq and processed data reported in this paper is GEO: GSE143295.

## Supporting Figure Legends

**Figure S1.**
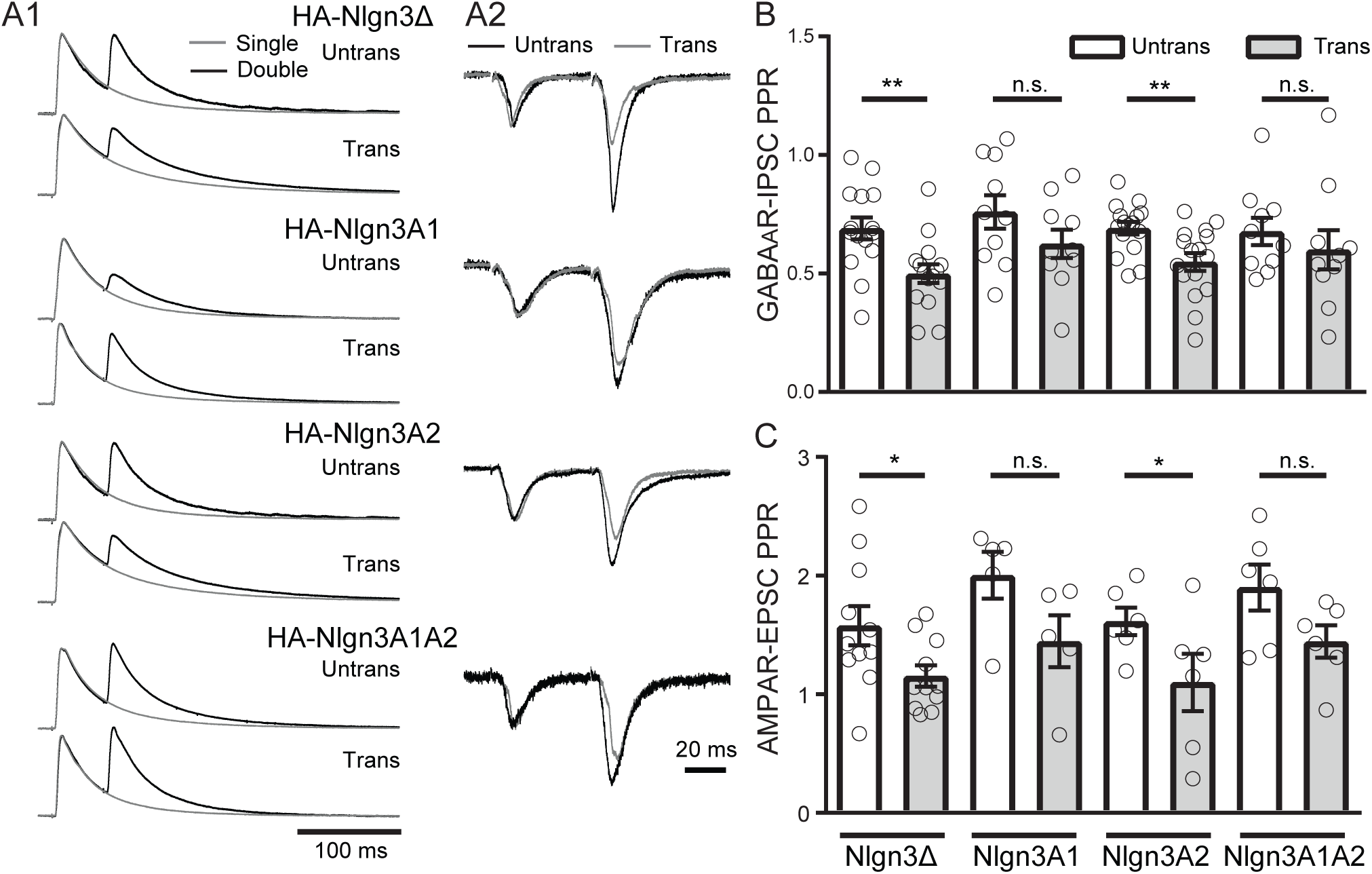
Nlgn3 splice isoforms differentially regulates presynaptic release probability. Effect of Nlgn3 splice isoform overexpression on PPR in hippocampal CA1 pyramidal cells. (A) Normalized sample traces of PPR of GABA_A_R-IPSC (left, A1) and AMPAR-EPSC (right, A2). IPSCs and EPSCs are normalized to the first amplitude. Because the first GABA_A_R-IPSC overlaps with the second IPSC, to accurately measure the amplitude of the second IPSC, we ‘cancelled’ the first IPSC by subtracting the traces receiving a single pulse (gray) from those receiving a paired pulse (black), both normalized to the first response. (B, C) Summary of effect of Nlgn transfection on IPSC- (B) and EPSC-PPR (C). Each bar represents the average of ratios obtained from multiple pairs of transfected and untransfected neighboring neurons. Number of cell pairs: Nlgn3Δ (GABA_A_R-IPSCs/AMPAR-EPSCs: 15/11); Nlgn3A1 (10/5); Nlgn3A2 (15/6); Nlgn3A1A2 (10/6). ** p < 0.01, * p < 0.05, ns, not significant. Student t-test.

**Figure S2.**
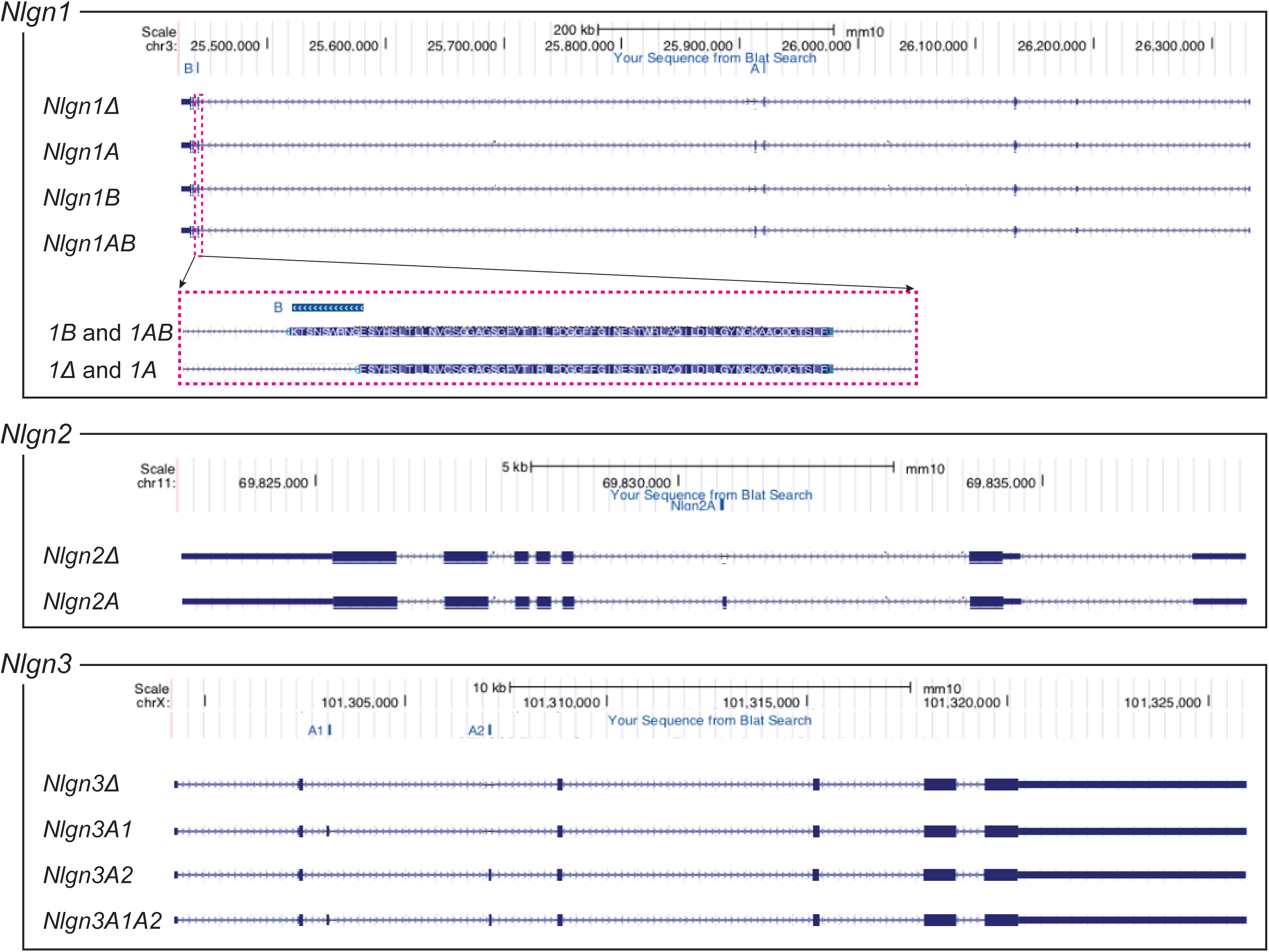
UCSC browser tracks of *Nlgn1, 2* and *3*. 10 *Nlgn* transcripts were manually curated.

